# Attention updates the perceived position of moving objects

**DOI:** 10.1101/543215

**Authors:** Ryohei Nakayama, Alex O. Holcombe

**Affiliations:** School of Psychology, The University of Sydney, NSW 2006, Australia; Center for Information and Neural Networks (CiNet), National Institute of Information and Communications Technology, 1-4 Yamadaoka, Suita City, Osaka, 565-0871, Japan

## Abstract

The information used by conscious perception differs from that which drives certain actions. An important example involves an object’s internal texture motion. The motion causes a perceptual shift that can accumulate over seconds into a large illusion, but targeting of the grating for a saccade (a rapid eye movement) is not affected by this illusion. One possible explanation is that rather than saccade targeting having privileged access to the correct position, the shift of attention thought to precede saccades resets the accumulated illusory position shift to zero. In support of this alternative to a dissociation between perception and action, we found that the accumulation of illusory position shift can be reset by transients near the moving object, creating an impression of the object returning to near its actual position. Repetitive color changes of the object also resulted in reset of the accumulation, less so when attention to the object was reduced by a concurrent digit identification task. Finally, judgments of the object’s positions around the time of saccade onset reflected the veridical rather than the illusory position. These results indicate that attentional shifts, including the shift preceding saccades, can update the perceived position of moving objects and mediate the previously reported dissociation between conscious perception and saccades.

## Introduction

Visual motion signals can strongly affect the perception of position. For example, texture motion within a stationary object shifts that object’s perceived position in the direction of the texture motion [1, 2]. Interestingly, if such an object moves in a trajectory orthogonal to its texture motion, the illusory position shift can steadily accumulate over seconds, giving a sensation of a trajectory oriented between the directions of texture and object motions. This is referred to as the infinite regress, curveball, or double-drift illusion [3, 4].

Lisi & Cavanagh [5] reported that when participants are asked to move their eyes (saccade) to a double-drifting object, their eyes land in approximately the veridical position rather than the illusory shifted position. Lisi & Cavanagh concluded that the information used by conscious perception is different from that which drives the saccadic eye movement. But the present results suggest that the attention shift thought to precede saccades resets the accumulated position shift to zero, explaining why saccades are made to the veridical object location, and highlighting a new role for attention.

Our first experiment shows that stimulus transients near the moving object can reset the accumulation of illusory position shift, creating an impression of the object returning to its actual position. Our second experiment provides evidence that repetitive color changes to the object also result in reset of the accumulation. The illusion was restored when attention to the object was reduced by a concurrent digit identification task. These results are consistent with the possibility that attention can reset accumulated position shifts.

In our third experiment, participants were asked to saccade to the moving object at the time of a signal (disappearance of the fixation point). As Lisi & Cavanagh found, the eyes landed close to the veridical position of the object. However, we also presented a probe object five hundred milliseconds after the saccade began. Participants judged the position of the probe relative to the moving object, and we aggregated the judgments across trials to reconstruct the average perceived position of the moving object after saccade initiation. The results indicate that the position shift accumulation had been reset. This finding suggests that a shift of attention toward the moving object updates a position estimate, which resets illusory position shift.

## Results

### Experiment 1

We characterized the trajectory participants perceive with a double-drift stimulus, and showed that irrelevant transients cause reset of the accumulation. We used objects consisting of gratings that were circularly windowed with a tapered edge. Two objects moved diagonally (Figure 1A), down the screen and outward from fixation, while the vertical gratings within them were either stationary within the circular envelopes (single-drifting objects) or drifted inward (double-drifting objects; See Movies S1-S3). During each single-drift trial and in half of the double-drift trials, white squares were flashed on both sides of each of the two moving objects for one movie frame, which occurred at a random time between 0.7-1.1 sec after stimulus motion onset. The moving objects disappeared 1.8 sec after onset. Participants maintained fixation throughout (monitored by an eye tracker) and drew the object trajectories they perceived with a computer mouse. All trials containing saccades in a period from 0.7 to 1.5 sec were removed from subsequent analyses.

**Figure 1.**
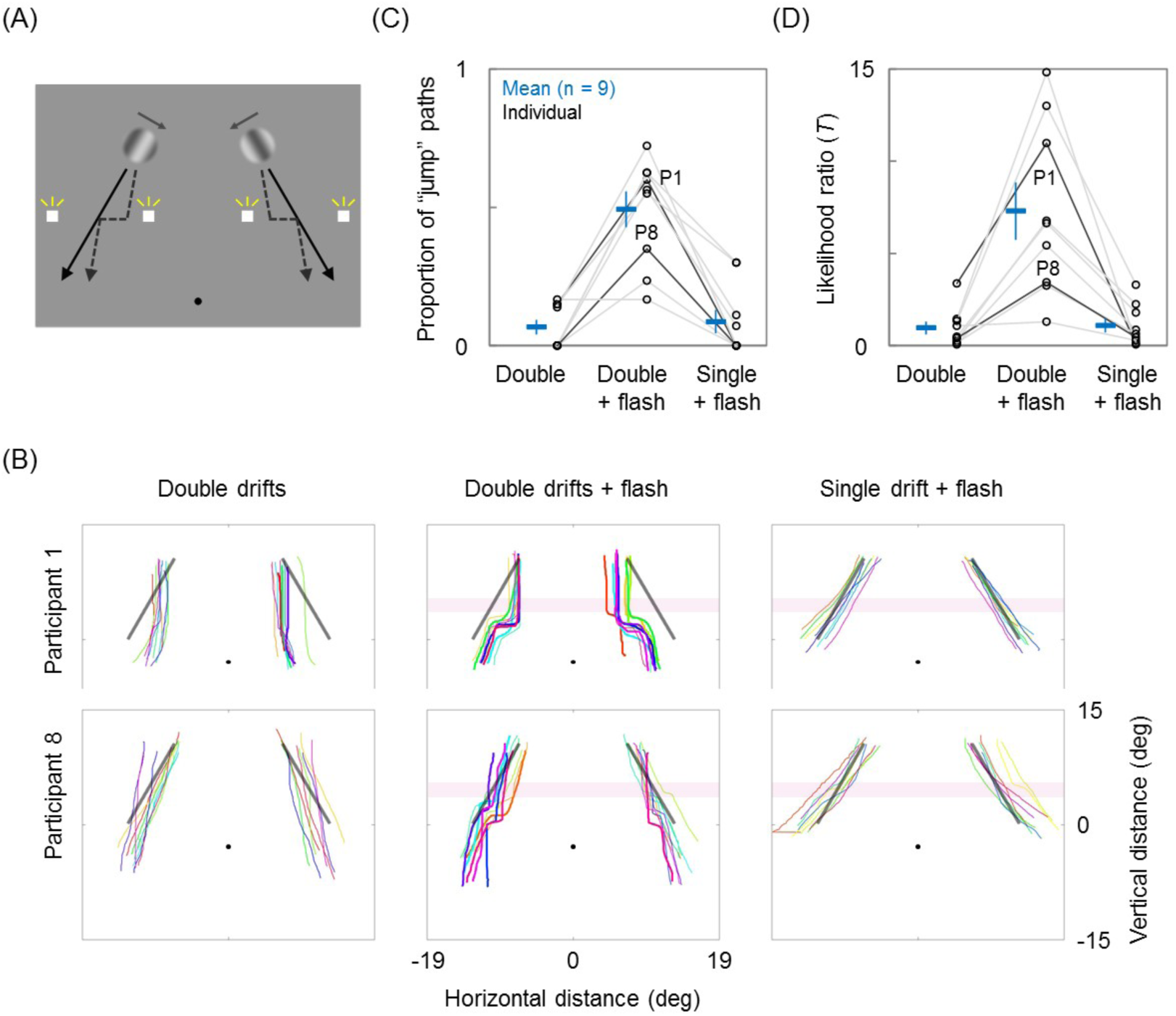
Schematic stimulus display and results of Experiment 1. (A) Two circular objects moved diagonally for 1.8 sec: double-drifting (with orthogonal drift of the internal pattern) or single-drifting (no drift of the internal pattern) objects (see Movie S1). Solid arrows (not shown during experiment) indicate the physical motion directions of envelope and grating for the double-drifting objects. White squares were flashed for one screen frame between 0.7-1.1 sec after the motion began, except there was no flash in half of the double-drift trials (Movie S2). Dashed arrows indicate possible perceived trajectories of double-drifting objects with flashes (Movie S3). (B) Object trajectories sketched by two representative participants (P1, P8). Rows correspond to participants and columns correspond to stimulus conditions. Display colors correspond to trial numbers and bold curves denote paths classified as jumps by maximum likelihood. Black diagonal lines indicate physical trajectories and pink-shaded bands indicate spatial locations of flash presentations. (C) Blue horizontal bars show average proportion of perceived trajectories classified as jumps (circles for individuals). (D) Blue horizontal bars show average likelihood ratio or *T* (circles for individuals). Error bars represent ±1 SE across participants.

Figure 1B shows the trajectories reported for double-drifting objects (left column), double-drifting objects accompanied by flashes (middle column), and single-drifting objects accompanied by flashes (right column). Pink-shaded bands indicate intervals wherein flashes were presented. It can be seen that the internal grating motion of the double-drifting objects biased the trajectories perceived toward fixation. The appearance of the flashes (middle column) frequently resulted in a sharp trajectory change, sometimes described by participants as involving a jump, toward the true position of the object. On the other hand, the single-drifting objects continued relatively unchanged. In summary, the luminance flash frequently reset the accumulation of illusory position shift induced by double drifts.

The variance of the trajectories drawn for different trials was non-trivial, even within a participant (see Figure S1 panels for the trajectories by all participants). To quantitatively and reproducibly assess the proportion of trials in which the drawings indicated a trajectory change, we classified them as “jump” and “no jump” trials with a likelihood-ratio test [6]: a statistical comparison of models with versus without a horizontal jump in a linear or curved course of stimulus trajectories. Specifically, a quadratic curve (*x* = *ay*^2^ + *by* + *c*; *x* and *y* denote horizontal and vertical positions; *a*, *b*, and *c* are free parameters) both with and without two additional parameters specifying a horizontal jump (corresponding to the timing and the size of the jump) was fitted to each trajectory, by maximum likelihood assuming that the model’s errors are normally distributed.

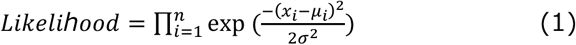

Here, μ and σ denote the model prediction of horizontal position (x) and standard deviation of horizontal positions, respectively. The fits of the curves were very good – on average the jump/no-jump model *r*^2^ = 0.956/0.948 (SE = 0.017/0.018) for double-drifting objects without flashes, 0.958/0.912 (SE = 0.008/0.019) with flashes, and 0.985/0.980 (SE = 0.005/0.006) for single-drifting objects with flashes (also see Figure S2 panels for the best-fit curves). The likelihood-ratio test was used to decide whether the jump model fit significantly better than the no-jump model. This test used the maximum likelihood for the jump and no-jump models and the following test statistic (*T)*, which follows a chi-squared distribution (degrees of freedom = 1; the timing parameter operates fully depending on the presence/absence of the size parameter).

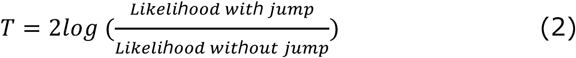

Perceived (drawn) trajectories were classified as “jump” paths if *T* values exceeded 3.84 (corresponding to *p*<.05), which are drawn in bold in Figure 1B. Figure 1C shows the proportion of the “jump” paths in each condition. Jumps were much more common for double-drifting objects with flashes compared to the other two conditions (one-way ANOVA *F*_(2,16)_ = 30.40, *p* < 0.001; between double-drifting objects with versus without flashes *t*_(16)_ = 6.91, *p* < 0.001; between double-versus single-drifting objects with flashes *t*_(16)_ = 6.59, *p* < 0.001), while there was no significant difference between double-drifting objects without flashes and single-drifting objects with flashes *t*_(16)_ = 0.32, *p* = 0.75. As an alternative analysis, instead of using the *T* value to classify the trials and comparing their proportion, we compared *T* values between conditions directly. Figure 1D plots average and individual *T* values. Similar to the classification-based analysis, the *T* value is significantly larger for double-drifting stimuli with flashes (one-way ANOVA *F*_(2,16)_ = 17.35, *p* < 0.001; *t*_(16)_ = 5.15, *p* < 0.001; *t*_(16)_ = 5.05, *p* < 0.001; *t*_(16)_ = 0.10, *p* = 0.92 in comparisons of the same order as above). The perceived jumps occurred around the time of the flashes (relative to the flashes’ time, −0.04 ± 0.31 sec, 95% CI), based on the curve fits after scaling the vertical distance of the sketched trajectory to that of the actual trajectory.

These results indicate that presentation of the luminance flashes reset the accumulated illusory position shift to near zero. Why do the flashes lead to a position reset? It might conceivably be mediated by the first-order motion energy of the flashes [7, 8], the fact that flashes can suppress certain percepts [9], or from the flashes capturing the participants’ attention.

### Experiment 2

If an attention shift is the cause of resets, an attentional load at fixation should reduce the occurrence of position resets, both without transients and in the presence of transients. To test this prediction, in Experiment 2 the participants were given an additional task of identifying digits embedded in a central stream of letters.

Specifically, we postulated that repetitive transients to the object, which are expected to regularly reset the accumulation, would result in the perceived path approximating to the physical tilt rather than the illusory shift. If attention is necessary for the occurrence of position resets, an attentional load would restore the illusory shift regardless of the presence of transients. A simple judgment of motion path tilt was utilized as it might be more robust to attentional demands than the previously used drawing task.

As a pre-test to allow nulling of the baseline illusion and thereby make any resets even more conspicuous, we estimated the subjective verticality of motion path orientation with a double-drifting object presented for 1.5 sec in the left or right visual field (Movie S4). After the participant indicated whether they perceived the path to be tilted to the left or to the right, for the next trial the physical path orientation was chosen by a staircase procedure designed to converge on the subjectively vertical orientation. The opposite texture motions, shifting the perceived path orientations in their directions, produced 33.5° (SE = 3.0°) difference in the subjective verticality on average (see Figure S3A).

In the main experiment, a double-drifting object moving at the path estimated as vertical in the pre-test underwent repetitive color changes between magenta and cyan at 1.5 Hz (Movie S5). The color changes consisted of a combined chromaticity and luminance modulation with one of three possible amplitudes for the luminance component – 0% corresponded to subjective equiluminance with the background (meaning only chromatic changes occurred) adjusted for each participant (see Methods). In the single task trials, participants were asked to judge whether the tilt of the object’ s path was to the left or to the right. In the dual task trials, tilt judgment was required only on trials where participants correctly identified two digits embedded at random times in a central stream of alphabetical letters. The central stream was presented both in the single and dual task trials and thus, the digit identification task reduced the participant’s attention to the moving stimulus without affecting stimulation of motion or luminance transient detectors.

Figure 2 shows the proportion of trials in which the true path tilt was reported. Chance reporting, 0.5, would correspond to the trajectory being perceived as vertical on average, as in the pre-test. We found that the correct tilt was perceived more often in the single task trials than in the dual task trials, and also more often with large luminance components (two-way ANOVA *F*_(1,7)_ = 14.71, *p* = 0.006 for task mode; *F*_(2,14)_ = 21.42, *p* < 0.001 for modulation strength; *F*_(2,14)_ = 0.59, *p* = 0.57 for interaction). This suggests that an attentional load reduces resets, preserving the illusory shift that might otherwise be eliminated by transients.

**Figure 2.**
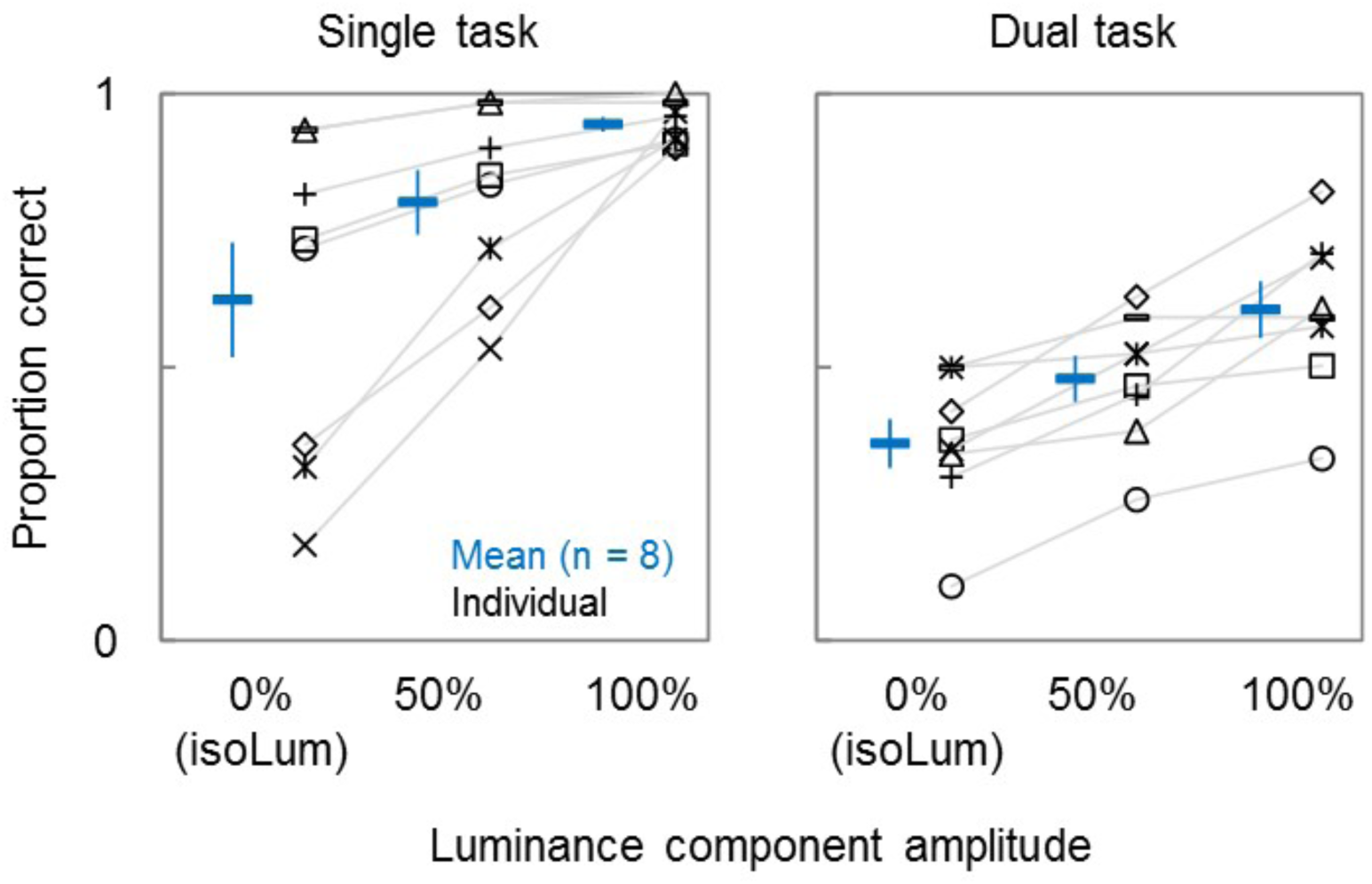
Results of Experiment 2. Blue horizontal bars show average proportion of trials in which the correct tilt was perceived for the subjectively-vertical (in the pre-test) trajectory (other markers show individual participants). Abscissa indicates amplitude of the luminance component of a combined chromaticity and luminance 1.5 Hz modulation. The stimulus display and primary task (path tilt judgment) were the same for the single and dual task trials. The dual task trials required participants to identify two digits from a central stream of letters, which substantially decreased the proportion of correct reports, suggesting it reduced the incidence of resets. Data is collapsed across path tilt and visual field conditions. Error bars represent ±1 SE across participants.

An alternative explanation for the dual-task finding is that discrimination thresholds increased so much that participants were unable to discriminate the orientation change caused by the resets. Resets likely changed the perceived tilt from approximately vertical to the physical tilt, which across participants averaged 16.7° (averaging the tilts of all the stimulus conditions; SD = 4.3°). But a control experiment indicates that the impairment to path discrimination thresholds by the dual-task demand was only about 1.5° (SD = 1.9°), and not significantly different from zero in three out of four stimulus conditions: left tilt in LVF *z* = 0.02, *p* =0.99; right tilt in LVF *z* = 2.66, *p* = 0.01; left tilt in RVF *z* = 0.82, *p* = 0.41; right tilt in RVF *z* = 1.08, *p* = 0.28, too small to prevent perception of the orientation change caused by the resets.

Strikingly, at subjective isoluminance (0% luminance amplitude) in the dual task trials, participants reported the incorrect tilt significantly more often than chance (*z* = 3.10, *p* = 0.002). Why did participants so often report the incorrect tilt? If resets occasionally occurred even in the pre-test, as expected, then the pre-test would somewhat underestimate the orientation needed for subjective verticality for trials without a reset.

This is because participants likely perceived close to the correct tilt in the trials with resets, biasing the staircase toward the convergence at a smaller tilt. In the dual task, a reduction in resets due to the dual task results in more accumulation of position shifts than in the pre-test, driving the perceived orientation beyond subjective vertical, so the incorrect tilt is reported. This account implies that attention causes spontaneous resets which are reduced by an attentional load.

Together, these results indicate that it is a shift of attention toward a moving object that triggers the position reset, not the luminance change.

### Experiment 3

This experiment was designed to examine more directly how saccades interact with the accumulated position shift in perception. If the attention shift preceding saccades causes resets, veridical positions should be perceived around the time of saccades targeting double-drifting objects.

Participants were asked to make a saccade to a double-drifting object moving along the path found to be subjectively vertical in the pre-test. The moment to saccade was indicated to the participant by disappearance of a fixation point at 0.3, 0.6 or 0.9 sec after motion onset. The average saccade latency was 300 msec (SD = 25 msec). The object was removed from screen as soon as the gaze was detected to be over 2 deg from the fixation point, which occurred on average 25 msec (SD = 1 msec) after the saccade onset and the saccade duration was 60 msec (SD = 6 msec), i.e., the object was removed before saccades landed. A probe was flashed 500 msec after the saccade onset and participants judged whether it was offset to the left or to the right of where the object was removed. After the participant’s response, the horizontal offset was adjusted for the next trial according to a staircase to estimate the point of subjective equality (PSE). This PSE provided a measure of where the moving object was perceived to be around the time of the saccade.

As can be seen in Figure 3A, the experiment replicated the finding of Lisi and Cavanagh (2015) that the path orientation represented by the saccade landing points (dots and dashed lines) is approximately the same as the physical path orientation (black lines). The two do not differ significantly from each other (for right tilt *t*_(5)_ = 0.22, *p* = 0.83, BF_10_ = 0.38; for left tilt *t*_(5)_ = 0.43, *p* = 0.69, BF_10_ = 0.40) but do differ from the subjective verticality (for right tilt *z* = 8.17, *p* < 0.001; for left tilt *z* = 4.74, *p* < 0.001). Also as Lisi & Cavanagh found, for some participants the saccades fall short of the physical positions, but for others they do not (see Figure S4 for the results of all participants).

**Figure 3.**
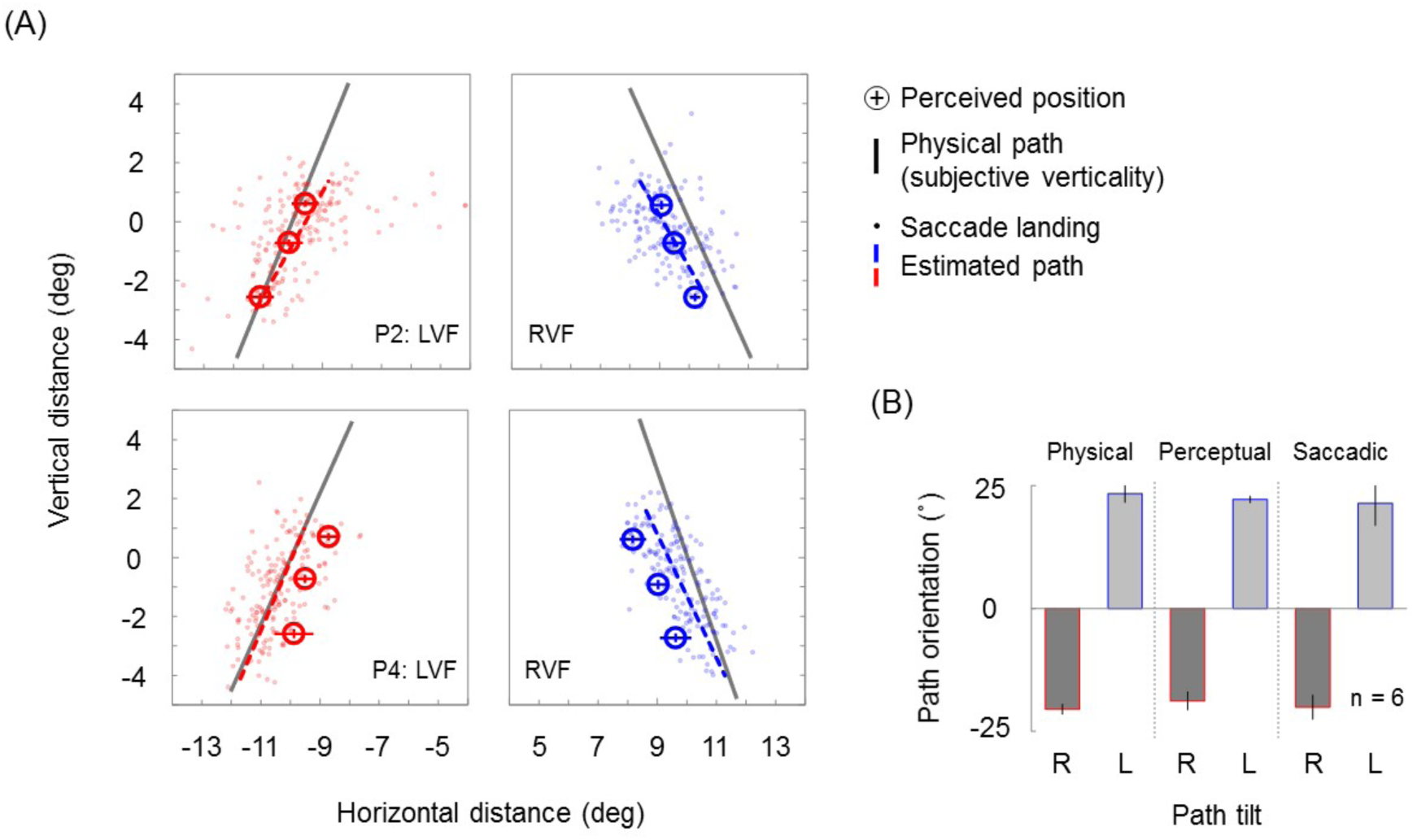
Results of Experiment 3. (A) Solid lines indicate physical path orientations (estimated without saccades in pre-test as subjectively vertical). Perceived positions (circles) and estimated motion paths (dashed lines) based on saccade landings (dots), for two of the six participants (P2, P4). The perceived positions were inferred from judgments about a probe presented soon after saccade initiation (signaled at one of three different times). Error bars represent 95% CI (horizontal bars derive 400 bootstraps). Dots indicate saccade landing points. Dashed lines are linear regressions of the saccade landings. Red/blue denotes left/right visual field (corresponding to right/left tilt in path orientation). (B) Average orientation of physical path (left), path estimated from perceived positions (middle) and path estimated from saccade landings (right). Error bars represent ±1 SE across participants.

Figure 3A also shows the perceived positions (circles). Just as for the saccade landing points, the orientation of the trajectory (linear regression) of the perceived positions is approximately equivalent to that of the physical trajectory (solid lines). These orientations are summarized in Figure 3B (physical versus perceptual panels) and are not significantly different from each other (for right tilt *t*_(5)_ = 0.89, *p* = 0.41, BF_10_ = 0.51; for left tilt *t*_(5)_ = 0.55, *p* = 0.61, BF_10_ = 0.42).

The resemblance of the perceived positions to the actual physical trajectory implies that they differ from the vertical orientation perceived during the pre-test where there was no saccade task, and indeed they are statistically significantly different from vertical (for right tilt *z* = 10.23, *p* < 0.001; for left tilt *z* = 32.60, *p* < 0.001).

These results indicate that the accumulation of illusory position shift is not reflected in perception around the time of the saccade. Combined with our previous results indicating that a shift of attention triggers the position reset, the results suggest that the attention shift preceding saccades resets the accumulated position shift to zero.

## Discussion

We found that an accumulated illusory position shift is often reset to approximately zero by a luminance or chromatic transient in a moving object. This reset, we suggest, is dependent on attention being attracted by a transient. In addition, the perceptual position shift was eliminated by the execution of a saccade to the moving object. These results are consistent with the proposition that a shift of attention, including those thought to precede saccades [10–12], can reset the illusory accumulation of the double-drift illusion.

Lisi & Cavanagh (2015) found that saccades to a double-drifting object were unaffected by the illusion. They suggested that this constituted the first clear and large difference between perceived position and saccade targeting, providing critical evidence for the long-running debate over how perception and action differ in their use of sensory signals. However, the present study suggests that the reason saccades are made to the correct position is best explained by the associated allocation of attention [13] rather than a dissociation between perception and action. This also explains jump percepts similar to those found here and caused by a gaze shift to a double-drifting object as described in the previous report of the double-drift illusion [3].

The present study is consistent with the possibility that rather than saccades and other actions having privileged access to the correct position, perceptual and oculomotor systems rely on the same position information [14]. Several previous studies have investigated stationary objects with internally-drifting texture, which elicits a small position shift and relatively little accumulation [1,2,15,16]. One study [17], while showing that the size of the shift varies with the stimulus used to compare position to, specifically found that it is *smaller* if compared to flashed lines than if judged by actions, specifically pointing and saccades. We suggest that this result may reflect the flashed lines resetting the position shift close to zero, whereas the actions are guided somewhat by information after the attention shift, during which position shifts may accumulate [15, 16].

While an eyetracker was used to exclude trials with saccades in Experiment 1, smaller eye movements, termed micro-saccades, may be a concern as abrupt stimulus changes induce micro-saccades [18]. However, as can be seen in Movie S6, the double-drift illusion still occurs when a fixation target in motion is pursued with the eyes. Smooth pursuit regularly contains small saccades to catch up to a moving target tracked but such small jitters in retinal image seem not to reset the accumulation of position shift.

The objects in the present experiments were likely attentionally tracked by the participants throughout most trials except the dual task trials, since the participants’ task was to draw or judge the tilt of their trajectories. Diluting attentional tracking resources among four such objects [19, 20] appears to have little effect on the double-drift illusion [21]. But the transients that accompany sudden visual changes have long been suggested to engage attention to a greater extent than typically occurs with top-down attention (e.g., [22]), and may even engage qualitatively different attentional functions than does top-down attention [23, 24], such as resetting the phase of ongoing oscillations [25]. However, because we found similar effects here with the saccade execution (or perhaps preparation) that involved no transients, we suspect that high intensity of top-down attention can cause reset.

Why would a greater intensity of attention reset the position perceived during the double-drift illusion? Attention to an object, even if its location is shifted, likely brings additional resources to bear for estimating object position, such as detection of prediction error (e.g., [26]) and activation of the primary visual cortex neurons stimulated by that object [27], with their greater spatial precision. To optimally use position and motion signals, greater position precision should reduce the contribution of the texture motion [28] that creates the illusion. Neurally, the primary visual cortex representations whose use improves position precision may themselves be shifted in the direction of object motion [29], so use of these representations should not completely eliminate the double-drift illusion. However, because the amount of shift in retinotopic areas is proportional to the size of the receptive fields [29], the use of early visual cortical areas should greatly reduce the size of the illusion relative to the use of later visual cortical areas. Thus an increase in attention can cause a perceptual reset of the accumulated position shift.

## Materials and Methods

### Participants

Nine observers (4 female, 1 author) participated in Experiment 1. Eight observers (6 female, 1 author) participated in Experiment 2. Six observers (4 female, 1 author) participated in Experiment 3. All participants had normal/corrected-to-normal vision. All experiments were conducted in accordance with the Declaration of Helsinki (2003) and were approved by the ethics committee of the University of Sydney. All participants provided written informed consent.

### Apparatus

Images were displayed on a gamma-corrected 22-inch CRT monitor (1280×1024 pixel) with a frame rate of 75 Hz. The CRT pixel resolution was 1.8 min/pixel at a viewing distance of 57 cm. Mean luminance was 45.8 cd/m^2^ and CIE chromaticity was (0.61, 0.35) in R, (0.29, 0.62) in G, and (0.15, 0.07) in B channels, respectively. Only in Experiment 2, where another CRT monitor was used, mean luminance was 74.8 cd/m^2^ and CIE chromaticity was (0.60, 0.33) in R, (0.29, 0.60) in G, and (0.16, 0.07) in B channels, respectively. Eye movements of the right eye were monitored at a sampling rate of 1000 Hz (EyeLink 1000 version 4.56; SR Research Ltd, Mississauga, Ontario, Canada).

### Stimuli

Visual stimuli were circular objects (1.8 deg in diameter) consisting of sine-wave grating texture (0.8 cycle/deg) enveloped by blurry edges (0.3 deg in cosine gradient). In Experiment 1, a pair of circular objects moved for 1.8 sec at 6.7 deg/sec diagonally (diagonally down and outward at an angle of 30° from straight down) in left and right upper visual fields respectively. Their initial position was 15.5 deg away from a black fixation point (0.2 deg in diameter). The orientation of each grating was parallel to the motion path, its luminance contrast was 0.5, and its initial spatial phase was randomized. The grating of the double-drifting objects drifted inward at 3.0 deg/sec. For the single-drifting object, it was stationary relative to the envelope.

In Experiment 2, a circular object moved for 1.5 sec at 6.7 deg/sec in either left or right visual field. The center of the motion path is level with but horizontally 10 deg away from a fixation point and the path orientation was variable across trials. The grating orientation was always parallel to the path orientation, its luminance contrast was 0.3, and its initial spatial phase was randomized. The grating drifted either inward or outward at 3.6 deg/sec. In Experiment 3, a circular object same as Experiment 2 was used except that the grating drifted only inward.

### Procedure

All experiments were conducted in a dark room.

#### Experiment 1

White squares (0.7 x 0.7 deg) were presented for 13.3 msec (one movie frame) on both sides of each of the two moving objects with horizontal offsets of 5.4 deg from them, which was randomly timed between 0.7 and 1.1 sec after the start of each trial. In each trial, participants maintained gaze on the fixation point throughout the moving object presentation, after the objects disappeared, traced the object trajectories by manually dragging a computer mouse. Participants had the option to re-draw their response as many times as they liked before confirming their response. Ten trials (except for Participant 9, who performed twenty trials) per stimulus condition were randomly interleaved. We removed trials that contained blinks or saccades (identified as eye movements over 30 deg/sec or 8000 deg/sec2) in a period from 0.7 to 1.5 sec after the start of each trial. On average, 81% (SD = 15%) of trials were used in subsequent analyses. The proportion of jump trials for each condition for each participant was analyzed with a one-way repeated-measures ANOVA.

#### Experiment 2

In the pre-test, participants judged the left/right tilt of the motion path after the presentation of a moving object in each trial. The physical orientation of the path was adjusted by the staircase 1 up 1 down rule with a step of 8 deg, which targets a 50% proportion of “right” tilt responses. A hundred trials per stimulus condition combining left/right tilt and visual field were randomly interleaved. For each condition, we estimated the point of subjective equality (PSE) as the vertical path corresponding to 50% correct (i.e., chance reporting of the tilt), calculated by the maximum likelihood method.

In the main experiment, the presentation of a moving object was always at the perceptually vertical orientation estimated above and, besides that, the object envelope was defined by a chromaticity modulation that alternates between magenta and cyan at 1.5 Hz stepwise. The red intensity of the envelope was minimized for cyan or maximized for magenta. The green intensity was chosen from three luminance component conditions such that color changes between magenta and cyan were subjective equiluminance or contained half or full modulation in mean luminance. Note that the luminance contrast of the grating was maintained at 0.3 across the conditions. Subjective equiluminance was determined before the experiment started by means of flicker photometry [30] – participants minimized the subjective flicker by adjusting the green intensity of the objects, positioned at the centers of the trajectories in both sides, alternating between the magenta and cyan colors at a rate of 8 Hz.

From 0.2 sec before to 0.2 sec after the stimulus-presentation period (1.9 sec in total), a rapid-serial visual presentation (RSVP) display appeared concurrently in the center of the screen instead of the fixation point. We used the RSVP display in order to confine the observers’ attentional resources throughout stimulus presentation [31]. In the RSVP display, 12 capital alphabetical letters (excluding I, O, Q, Y, Z) were serially presented every 80 msec and separated by a blank interval of 80 msec (6.3 Hz). Each letter was drawn in Arial font in black and subtended 0.75 x 0.75 deg. Two letters were replaced by two digits chosen at random between 1 and 9. One of the two digits appeared somewhere within the 2nd-to-5th letter sequence, and the other digit appeared within the 7th-to-11th letter sequence.

Single and dual tasks were tested in separate blocks. In single task trials, participants viewed the display with a steady fixation on the central RSVP letters and judged the left/right tilt of the motion path. Participants were instructed to concentrate on the moving object while gazing at the central letters. In dual task trials, participants were first asked to identify the two digits in the central RSVP display. If they identified both digits by pressing buttons in the correct order, participants then judged the left/right tilt of the motion path. On average, participants were able to respond within a few seconds after the stimulus presentation. Auditory feedback was given on each task. Participants were instructed to keep digit identification performance as high as possible. In dual task trials, all participants received practice trials in advance and only trials in which participants correctly identified the central digits were used in subsequent analyses. The average proportion correct of digit identification was 92.6% (SD = 3.7%). Thirty trials per stimulus condition combining luminance component, left/right tilt and visual field were randomly interleaved in each task block. Dual task trials incorrect in digit identification were not counted in this number.

#### Experiment 3

The presentation of a (gray-scale) moving object was always at the perceptually vertical orientation estimated in the same way as in the pre-test of Experiment 2. Each trial started when participants maintained gaze on the fixation point for 1 sec. Participants were instructed to make a saccade to the moving object as soon as the fixation point disappeared. The fixation point disappeared at one out of three points in time: 0.3, 0.6 or 0.9 sec after the stimulus started to move. As soon as the gaze position was detected outside a circular area with 2 deg of radius around fixation, the object was removed from screen so that participants received no feedback about the accuracy of their saccades. Five hundred milliseconds after the removal of the object, a white rectangle (H0.3 x V1.8 deg) was presented for 250 msec horizontally offset from the object’s last position.

Participants judged the left/right offset of the rectangle. The physical offset was adjusted by the staircase 1 up 1 down rule with a step of 0.9 deg. Sixty trials per stimulus condition combining left/right visual field and saccade-signaling time were randomly interleaved. Gaze position was recorded at 1000 Hz and monitored online; trials in which participants shifted gaze or blinked before the disappearance of the fixation point and trials with saccades not between 0.1 and 0.6 seconds after the disappearance were discarded online (not counted).

We removed trials with saccade landing points within 4 deg of radius around fixation. The average proportion of valid trials was 97.1% (SD = 2.3%). For each condition, we estimated the point of subjective equality (PSE) as the object’s last position corresponding to 50% correct (i.e., chance reporting of the offset), calculated by the maximum likelihood method. Then for each participant we computed a linear regression of the perceived positions against the three different saccade-signaling times, where horizontal coordinates were estimated as PSEs and vertical coordinates were actual last positions averaged across trials. The regression slope is the inferred perceived trajectory.

To recover the trajectory of the saccade targeting from the widely-distributed landing points, we fitted a multivariate linear model with the horizontal and vertical saccade amplitudes as dependent variables for each participant [5]. The models included as linear predictors the horizontal and vertical coordinates of the object at the moment of saccade onset. Then we used the fitted model to predict saccade amplitudes for each of the positions along the object’s path. Finally, we computed a linear regression of the vertical on the horizontal predicted saccade amplitudes between trials and derived the angle of deviation from vertical based on the regression slope recovered from saccades.

We tested the null hypothesis that the path orientation is equivalent between the physical and the inferred trajectories by the Bayesian paired samples t-test provided in the JASP statistical software [32]. BF_10_ is the ratio of the likelihood of the data under the alternative hypothesis divided by that under the null hypothesis. Values smaller than one indicate the data favor the null hypothesis over the alternative hypothesis.

## Acknowledgments

This work was supported by JSPS KAKENHI Grant Numbers JP18J01398 to RN. This study was carried out when RN was a JSPS research fellow, a visiting researcher at the University of Sydney, and a visiting researcher at CiNet, NICT.

## Supporting information

**Figure S1.**
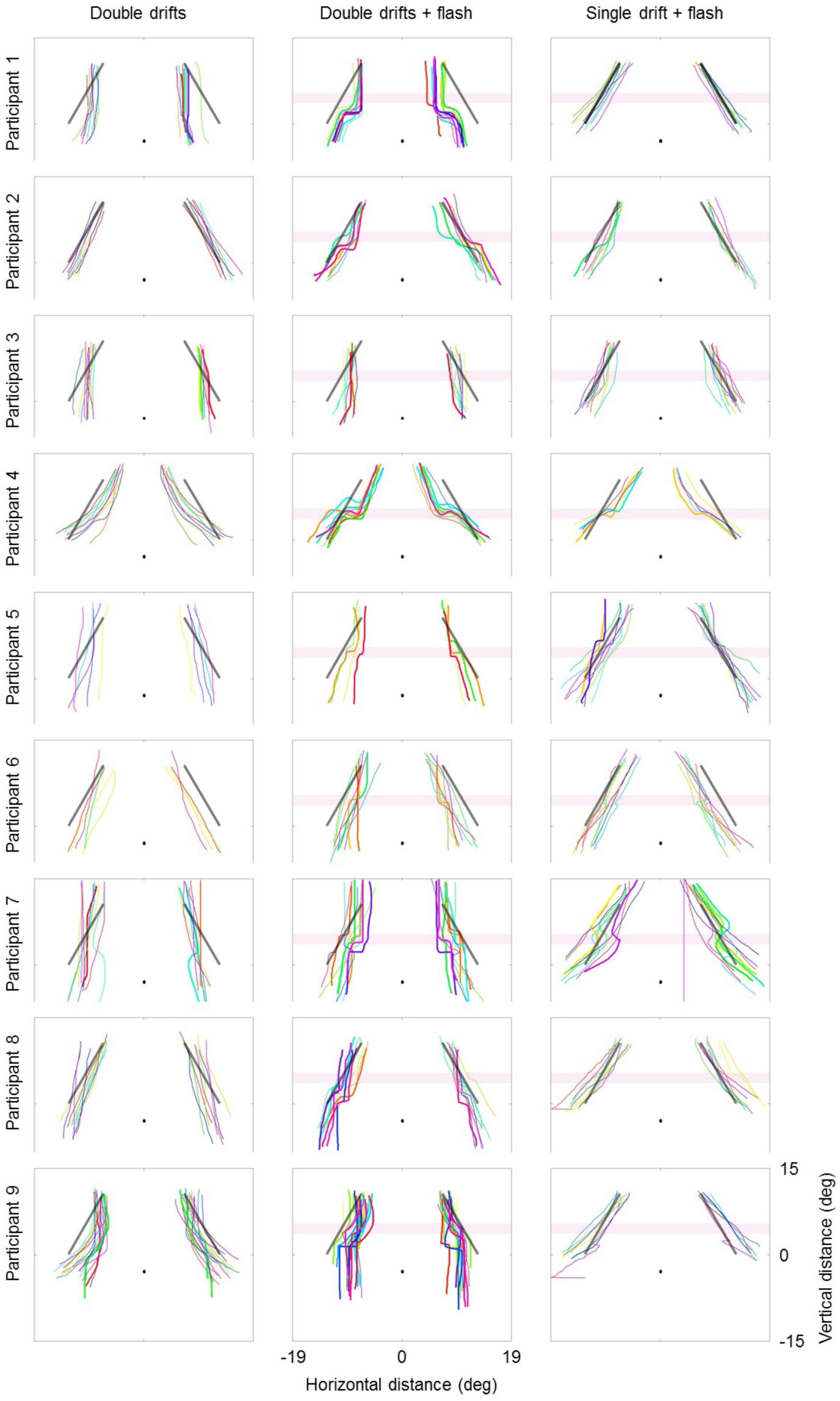
Object trajectories sketched by all participants in Experiment 1. View of the panels is same as Figure 1B.

**Figure S2.**
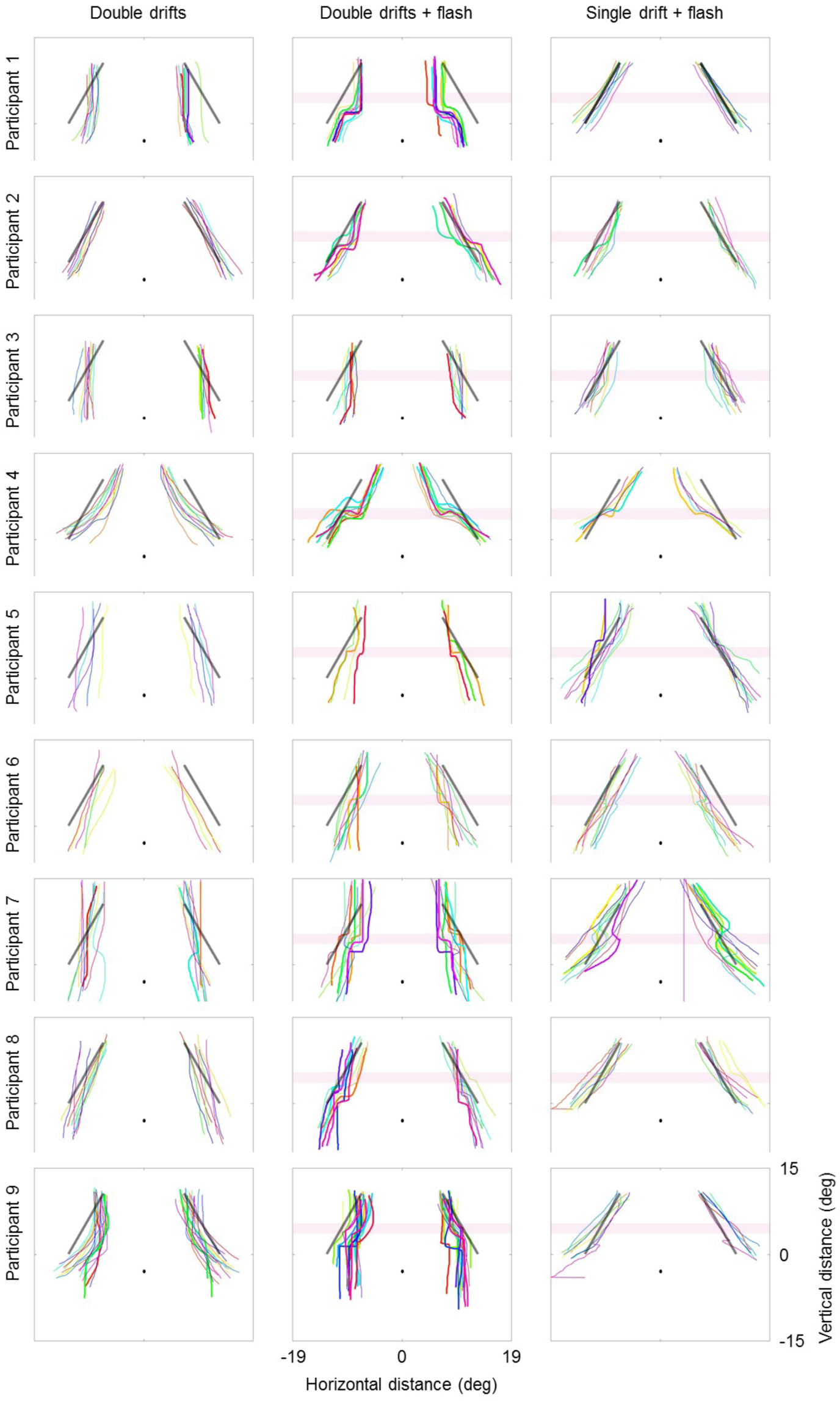
Best fits of object trajectories sketched by all participants in Experiment 1. View of the panels is same as Figure 1B.

**Figure S3.**
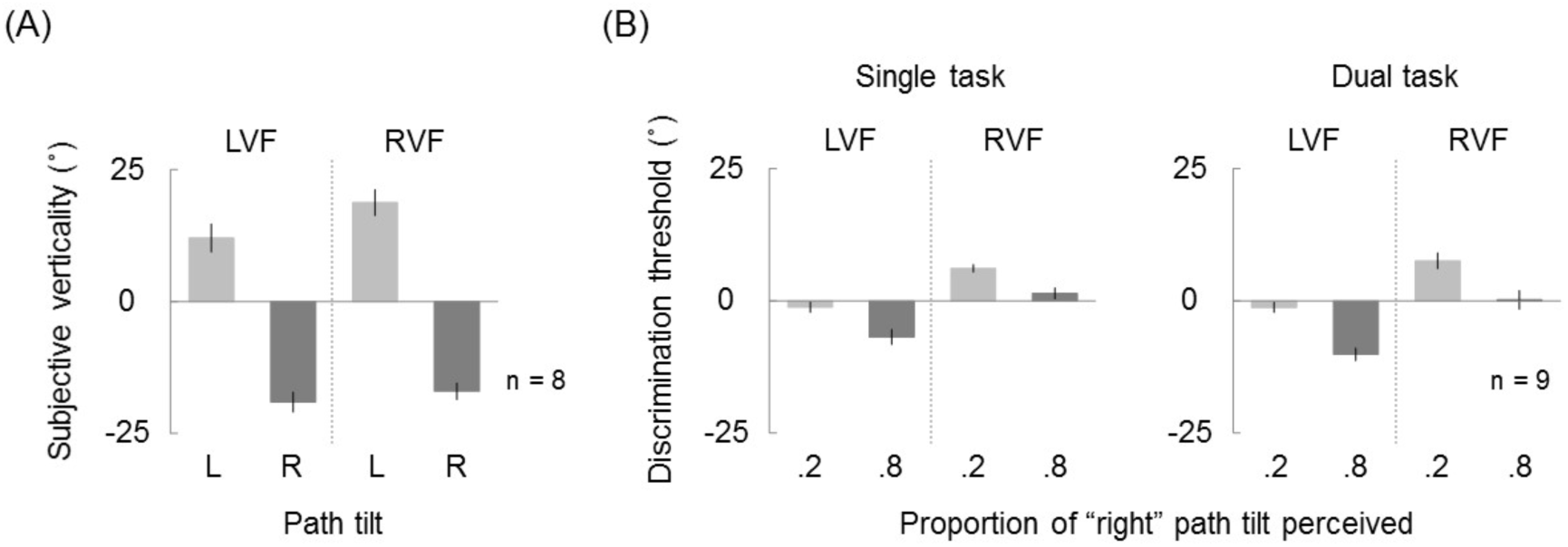
Results of the pre-test (A) and control experiment (B) for Experiment 2. (A) The path orientation, determined by pre-test, that was subjectively vertical (the orientation for which participants were equally likely to give the left/right path tilt response) for each combination of left/right path tilt and visual field. These orientations were used in the main experiment. (B) Thresholds for discriminating motion path tilt (mean of nine participants) at 20% and 80% accuracy levels, showing little effect of the dual task. Error bars represent ±1 SE across participants. In each trial, a single-drifting vertical grating object was presented in the same manner as the pre-test. Single task (tilt judgment only) and dual task (tilt judgment with digit identification same as the main experiment) were tested in separate blocks. The average proportion correct of digit identification was 90.0% (SD = 7.3%). Three-way ANOVA reveals a significant interaction between task and reporting rate conditions (*F*_(1,8)_ = 5.64, *p* = 0.045). Discrimination threshold at 80% in LVF is diminished by dual task (*t*_(8)_ = 2.66, *p* = 0.03, BF_10_ = 2.78) but not affected in the other conditions (at 20% in LVF *t*_(8)_ = 0.02, *p* = 0.99, BF_10_ = 0.32; at 20% in RVF *t*_(8)_ = 0.82, *p* = 0.44, BF_10_ = 0.42; at 80% in RVF *t*_(8)_ = 1.08, *p* = 0.31, BF_10_ = 0.51).

**Figure S4.**
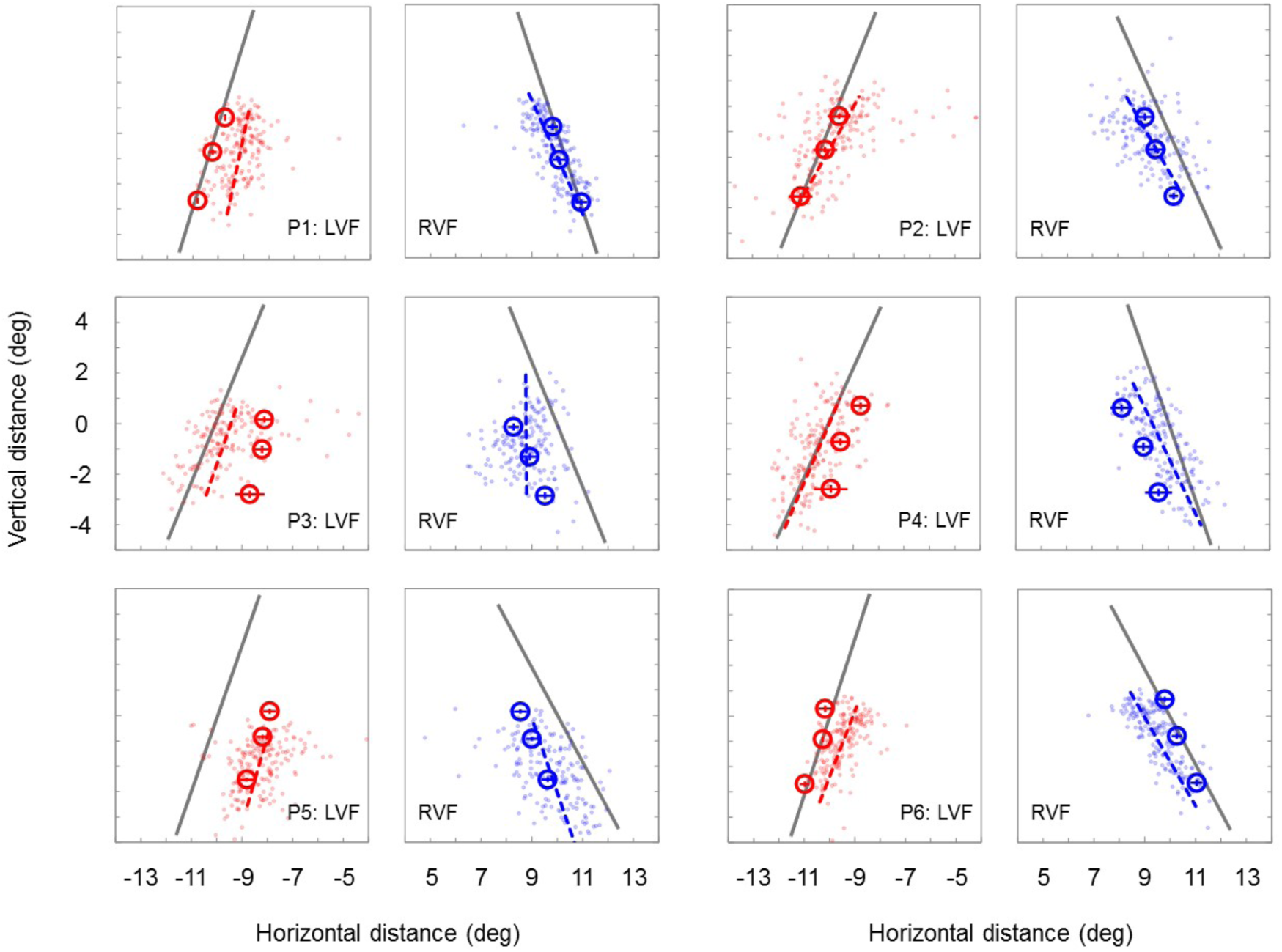
All participants’ results of Experiment 3. View of the panels is same as Figure 3A.

## Supplementary movies

View all videos in loop mode maintaining gaze at a fixation point. Repeated presentations seem to enhance the illusion.

Movie S1. Single-drifting objects accompanied by flashes. Diagonal trajectories will be perceived as they are mostly regardless of luminance flashes (control condition).

Movie S2. Double-drifting objects. Texture motion will accumulate over seconds into a large illusory position shift of objects, whose diagonal trajectories will be perceived as straight down or slightly slanted in periphery (double-drift illusion).

Movie S3. Double-drifting objects accompanied by flashes. Accumulated position shift will be reset to near zero by luminance flashes. If one tracks a separation between two moving objects, a flash presentation will apparently increase the separation, changing object trajectories.

Movie S4. An example of double-drifting objects used in the pre-test. Accumulated position shift will result in perception of the incorrect path tilt.

Movie S5. Accumulated position shift will be reset regularly by a combined color and luminance modulation, allowing observers to perceive the correct path tilt. However, if attention to the object is reduced by a second task at fixation (digit identification), the incorrect path tilt will be perceived despite the presence of transients.

Movie S6. Double-drift illusion still occurs during smooth pursuing with the eyes a fixation target moving in parallel with the object, showing that eye movement is not a direct cause of resets.

